# Sex differences in socioemotional behavior and changes in ventral hippocampal transcription across aging in C57Bl/6J mice

**DOI:** 10.1101/2023.05.01.538951

**Authors:** Nina E. Baumgartner, Mandy C. Biraud, Elizabeth K. Lucas

## Abstract

Socioemotional health is positively correlated with improved cognitive and physical aging. Despite known sex differences in socioemotional behaviors and trajectory of aging, the interactive effects between sex and aging on socioemotional outcomes are poorly understood. We performed the first comprehensive assessment of sex differences in socioemotional behaviors in C57Bl/6J mice across aging. Compared to males, females exhibited decreased anxiety-like behavior and social preference, but increased social recognition. With age, anxiety-like behavior, cued threat memory generalization, and social preference increased in both sexes. To investigate potential neural mechanisms underlying these behavioral changes, we analyzed transcriptional neuropathology markers in ventral hippocampus and found age-related changes in genes related to activated microglia, angiogenesis, and cytokines. Sex differences emerged in timing, direction, and magnitude of these changes, independent of reproductive senescence in aged females. Interestingly, female-specific upregulation of autophagy-related genes correlated with age-related behavioral changes selectively in females. These novel findings reveal critical sex differences in trajectories of ventral hippocampal aging that may contribute to sex- and age-related differences in socioemotional outcomes.

## 1. Introduction

Among the wide variety of cognitive and physical changes that occur with aging, socioemotional wellbeing emerges as a crucial factor in mitigating negative health outcomes in elderly populations (Charles and Carstensen 2010). Clinical literature has consistently shown a positive relationship between socioemotional health and improved cognitive (Seeman et al. 2001) and physical (Gill et al. 1997) outcomes in older adults. Although sex differences are observed across multiple species in both the trajectory of aging (Lemaitre et al. 2020, Hagg and Jylhava 2021) and the risk of socioemotional dysregulation (Kessler et al. 1994, Weissman et al. 1996, Gater et al. 1998), relatively little is still known regarding the interactions between age and sex on socioemotional behavior.

Preclinical rodent behavioral testing paradigms currently provide the greatest insight into the underlying biology of these behaviors (Rodgers et al. 1997, Stanley and Adolphs 2013). Of the studies that have explored sex differences in socioemotional behavior in rodents, results seem to vary greatly based on age, species, and genetic strain of subjects, as well as based on the types of behavioral tests used (Frick et al. 2000, Perkins et al. 2016, Domonkos et al. 2017, Hernandez et al. 2020). Sex differences have been reported in anxiety-like behavior (Kokras and Dalla 2014), social investigation (Tejada and Rissman 2012), and emotional memory (Bauer 2023), among other behaviors. Across aging, rodent models demonstrate increases in anxiety-like behavior (Darwish et al. 2001, Boguszewski and Zagrodzka 2002, Narita et al. 2006, Turner et al. 2012, Stanojlovic et al. 2019, Li et al. 2020, Hirano et al. 2021, Yanai and Endo 2021), decreases in social behaviors (Guan and Dluzen 1994, Boguszewski and Zagrodzka 2002, Salchner et al. 2004, Hunt et al. 2011, Perkins et al. 2016, Shoji et al. 2016, Gerasimenko et al. 2020), and impairments in emotional memory processes (Stoehr and Wenk 1995, Oler and Markus 1998, Houston et al. 1999, Doyere et al. 2000, Corcoran et al. 2002, Liu et al. 2003, Feiro and Gould 2005, Gemma et al. 2005, Gould and Feiro 2005, Moyer and Brown 2006, Fukushima et al. 2008, Kaczorowski and Disterhoft 2009, Peleg et al. 2010, Villeda et al. 2011, Shoji et al. 2016, Aziz et al. 2019, Shoji and Miyakawa 2019, Ehlers et al. 2020, Yanai and Endo 2021, Hernandez et al. 2022). Despite these clear effects of age and sex on socioemotional behaviors, shockingly few studies have directly compared these behaviors in males and females across aging. A notable connection between these behaviors is their regulation by the ventral hippocampus (Bannerman et al. 2004, Fanselow and Dong 2010), a brain region known to be sensitive to both age and sex (Wang et al. 2019, Williams et al. 2020, Porcher et al. 2021, Hodges et al. 2022).

Understanding how these sex differences in socioemotional behavior change across the lifespan and how that relates to ventral hippocampal function is therefore of critical clinical importance, especially considering that women represent the largest proportion of the aging population (Hagg and Jylhava 2021).

Here, we sought to establish the first comprehensive assessment of sex differences in socioemotional behavior across aging in C57Bl/6J mice. Male and female mice aged 4-, 10-, or 18-months, corresponding to young, middle-aged, and aged human populations, respectively (Flurkey et al. 2007), were subjected to a battery of behavioral tests related to socioemotional behavior. Female estrous cycles were monitored throughout the duration of the experiment in order to classify cycling regularity as a measure of reproductive senescence in females across aging. Following behavioral testing, ventral hippocampal tissue was analyzed for mRNA quantification using the NanoString Neuropathology panel to assess expression of 760 transcripts across aging in males and females. Our unique approach combining comprehensive behavioral testing and transcriptional analyses in the same subjects provides novel insight into sex- and aging-related changes in socioemotional behaviors and reveals critical sex differences in the trajectory of ventral hippocampal aging.

## 2. Materials and method

### 2.1 Subjects

Male and female C57BL/6J mice (#00064, Jackson Laboratories, Bar Harbor, ME) were bred in-house for the purposes of this study. Gonadally-intact mice at 4 (male, n=14; female, n=13), 10 (male, n=11; female, n=12), or 18 (male, n=12; female, n=13) months of age were used for experimental procedures. Mice were housed 2-4 per cage with corncob bedding and *ad libitum* access to standard chow (LabDiet Rodent 5001) and water in an environmentally controlled husbandry room maintained on a 12h:12h light:dark cycle (lights on at 08:00 AM). Male and female mice of each age group were tested in separate cohorts to avoid potential confounds of sex pheromones on behavioral assays. All experiments were approved in advance by the Institutional Animal Care and Use Committee at North Carolina State University and conducted in accordance with the National Institutes of Health *Guide for the Care and Use of Laboratory Animals*.

### 2.2 Estrous Cycle Categorization

To categorize the reproductive cycle, mice received vaginal (female) or sham (male) lavages daily between 8:00-9:00AM, starting 8 days prior to behavioral testing and continuing to euthanasia. Cells were stained using hematoxylin and eosin, and estrous stages were categorized using standard cytological methods (Cora et al. 2015). Briefly, proestrus was determined by the predominance of nucleated epithelial cells, estrus by the predominance of cornified epithelial cells, and diestrus by the presence of leukocytes. Estrous cycle regularity was categorized as regular, irregular, or non-cycling. Irregular cycling was defined as ≥5 days in the same cycle stage, and non-cycling was defined as a total absence of transition of a full cycle (estrus to estrus) for the duration of the experiment (19 days).

### 2.3 Behavioral Testing

Behavioral testing commenced at 9:00AM, and all mice were handled and tested by the same researcher (M.C.B.). Male and female mice were tested in separate cohorts, and equipment was broken down and thoroughly cleaned between cohorts. Mice were habituated to an airlock adjacent the behavioral testing room 30 mins prior to each behavioral test. All mice went through an identical testing sequence: Days 1-3, handling; Day 4, elevated plus maze; Day 5, open field test; Day 6, novel object recognition and object location; Day 7, social interaction and social recognition; Day 8, auditory threat conditioning; Day 9, cued and contextual threat memory recall. Behavioral data were analyzed blind to age and sex.

#### 2.3.1 Elevated Plus Maze

The plus maze consisted of two open arms (30 cm) and two closed arms (30 cm with 15 cm walls) raised 40 cm from the table surface (Panlab, Barcelona, Spain). The arena was indirectly lit with the center and open arms at 120 lux. Each mouse was placed in the center and allowed to explore the maze for 5 minutes. Behavior was recorded with an overhead camera and analyzed offline with AnyMaze software (Stoelting, Wood Dale, IL). Variables of interest included open arm time and percent open arm entries. Percent open arm entries was calculated as the number of open arm entries divided by the summed number of open and closed arm entries x100.

#### 2.3.2 Open Field Test

The apparatus consisted of an opaque gray arena (45 x 45 x 40 cm; Panlab) indirectly lit at 85 lux. Mice were placed in the corner of the arena and allowed to explore for 10 min. This test was repeated three times, with mice returned to a holding cage for 20 min between each test. Behavior was recorded with an overhead camera and analyzed offline with AnyMaze software. The center zone was set as the inner 225 cm^2^ of the area. Distance traveled (m) and center time (s) were analyzed and averaged across the three trials.

#### 2.3.3 Object Location and Object Recognition Tests

Three suction toys of similar size but distinct shape and color were used as objects, and objects were placed 11 cm from the corners of the open field area. For baseline exploration, two objects were placed in opposite corners. Mice were placed in a corner of the arena not containing an object and allowed to explore for 10 min before returning to a holding cage for a 20 min break. For the object location test, one of the objects was moved to a different corner of the arena, and mice were placed back into the arena for 10 min before returning to a holding cage for a 20 min break. For the novel object recognition test, the object that remained unmoved in the location test was replaced with a novel object. Mice were placed into the arena for 10 mins before returning to their home cage. Each test was recorded with an overhead camera, and behavior was analyzed offline using AnyMaze software to manually key interaction time (s) with each object. While a pilot group demonstrated no baseline preference for the objects used, experimental mice displayed a clear preference for one of the objects and thus confounded the data. Therefore, these results are not reported.

#### 2.3.4 Social Interaction and Social Recognition Tests

The three-chamber social interaction arena consisted of three chambers (20 x 42 x 22 cm) with transparent walls and removable doors separating the center chamber from the two outer chambers (Panlab). Mice were habituated in the center chamber for 5 mins. For the social interaction test, two identical grid enclosures containing either a novel age- and sex-matched conspecific mouse (Stranger 1) or a small rubber duck (Object) were placed in the outer chambers. The doors separating the inner and outer chambers were raised, and the test mouse was allowed to explore the entire apparatus for 10 min. The test mouse was then returned to a holding cage for a 5 min break, during which time the object was replaced with a novel age- and sex-matched conspecific (Stranger 2). For the social recognition test, each test mouse was placed in the center chamber with the doors to the outer chambers raised and allowed to explore the entire apparatus for 10 mins. Position of the Object, Stranger 1, and Stranger 2 were counterbalanced between animals. Each session was recorded with an overhead camera, and behavior was analyzed offline using AnyMaze software to manually key interactions with the object and novel conspecifics. Results for social interaction are expressed as social preference, calculated as a ratio of the time (s) spent investigating Stranger 1 versus the time (s) spent investigating the Object. Results for social recognition are calculated as a ratio of the time (s) spent investigating Stranger 2 versus the time (s) spent investigating Stranger 1. One cage of mice from each age cohort per sex were used as the novel conspecifics and were therefore excluded from subsequent behavioral testing and transcriptional profiling.

#### 2.3.5 Auditory Threat Conditioning and Recall

Cued threat conditioning, cued memory recall, and contextual memory recall were conducted in Habitest modular operant chambers (17.78 cm x 17.78cm x 30.48cm) housed in sound-attenuating cubicles (Coulbourn, Holliston, MA). The conditioning context consisted of two clear Plexiglas walls and two stainless steel walls with an aluminum shock grid floor, and 70% ethanol was used for the odorant. A near infrared camera was mounted behind each operant chamber, and FreezeFrame4 software (Actimetrics) was used for automated stimulus delivery and video recording. Auditory threat conditioning consisted of a 240 s baseline period followed by three co-terminating presentations of the conditioned stimulus (CS; 20 s, 2kHz, 80 dB pure tone) with the unconditioned stimulus (US; 2 s, 0.7 mA foot shock), with 120 s inter-trial intervals (Lucas et al. 2014). Thirty seconds after the final stimulus presentation, mice were removed from the operant chamber and placed into a new home cage. Cued and contextual recall were assessed 24 hrs later. For cued recall, the walls and the floor of the chamber were covered in white and blue striped inserts, and isopropanol was used as the odorant. Following a 180 s baseline period, 4 CSs were presented with 80 s inter-trial intervals. Mice were removed from the operant chamber and returned to their home cage for 2 hrs until contextual recall. Contextual recall occurred over 3 mins in the conditioning context in the absence of stimuli.

Freezing was analyzed as the conditioned response using Actimetrics FreezeFrame V4 software. Freezing thresholds were set for each mouse, determined by the highest movement index value representing no movement except for that required for respiration for at least 3 s (du Plessis et al. 2022). Freezing was measured during the baseline period and CS presentations during training and cued recall and across the entire 3 min trial for contextual recall. To control for the confound of group differences in baseline freezing, baseline freezing levels were subtracted from our cued and contextual recall analyses (Anagnostaras et al. 2010, Jacobs et al. 2010). For cued recall, pre-CS baseline freezing was subtracted from averaged CS freezing. For contextual recall, baseline freezing prior to conditioning was subtracted from total freezing across the 3 min trial.

### 2.4 Nanostring Sample Preparation and Analysis

Five days after threat memory recall, mice were deeply anesthetized with Avertin (500 mg/kg; Thermo Scientific, Waltham, MA) prior to decapitation. Brains were rapidly extracted and flash frozen with 2- methylbutane on dry ice prior to storage at -80°C. For 18-month-old females, the entire uterine horn was dissected and weighed at the time of euthanasia to determine uterine index, calculated as the weight of the uterine horn divided by total body weight. One 18-month female was excluded from uterine index analysis due to extreme ovarian swelling, likely due to presence of tumor(s) (Smith and Xu 2008).

Brains from a subset of animals from each cohort were selected for transcriptional profiling. Animals were chosen to represent all litters and cages that underwent all behavioral assays, and females from the 18-month group included both cycling and non-cycling animals. Brains were sectioned at a 500 μm, and the entire right ventral hippocampus was collected via 1 mm diameter micropunches at -2.8mm from Bregma using the Palkovits method (Palkovits 1983). Punches were stored at -80°C until RNA extraction. Tissue was homogenized in Trizol reagent followed by RNA isolation with the Qiagen RNeasy Mini Kit (Germantown, MD) following the manufacturer’s instructions. Isolated RNA was purified and concentrated with Amicon Ultra-0.5 Centifugal Filters (Millipore Sigma, Burlington, MA) prior to storage at -80°C.

The Nanostring Neuropathology panel (NanoString Technologies, Seattle, WA) was used for transcriptional profiling of 760 targets using highly sensitive outputs of mRNA transcript counts via color-coded reporter probe detection (Geiss et al. 2008). This technology has repeatedly been shown to achieve the accuracy of quantitative real-time PCR with enhanced sensitivity for detecting low- abundance transcripts without relying on amplification steps (Geiss et al. 2008, Malkov et al. 2009, Reis et al. 2011, Veldman-Jones et al. 2015). RNA was diluted to 20 ng/μL, and a total of 100 ng of RNA per sample was hybridized to the capture probesets at 65°C for 21 hrs. Hybridized samples were immediately processed on the Nanostring nCounter, which purifies and aligns samples onto the internal surface of the sample cartridge prior to barcode reading on the digital analyzer. Twelve samples were processed per cartridge, and a calibrator sample was used to normalize data between cartridges. Raw data were assessed for quality control parameters including binding density of barcodes measured, image quality for each well, assay efficiency, and linearity via comparisons to positive and negative controls for each well.

Raw transcript counts for each gene were normalized to the geometric mean of 10 housekeeping genes: *Aars, Asb7, Ccdc127, Cnot10, Csnk2a2, Fam104a, Lars, Mto1, Supt7l, Tada2b*. The expression of the individual housekeeping genes, as well as their geometric mean, was not found to differ across age or sex. Counts were further normalized to the calibrator sample on each cartridge to normalize for variability between cartridges. To determine differentially expressed genes, pairwise comparisons of normalized transcript counts were performed between ages within each sex. A threshold of Benjamini-Hochberg False Discovery Rate (FDR) adjusted p-value of < 0.1 was used to set statistical significance for differentially expressed genes (Benjamini et al. 2001).

Transcript data was further analyzed using the nSolver Pathway Analysis Tool, which groups genes into pre-assigned pathways based on putative gene function and reports a pathway score as the log2 fold change versus a reference group. For pathway comparisons examining the effects of aging within each sex, the younger age group for each sex was considered the reference group. For pathway comparisons examining the effect of reproductive senescence in 18-month females, cycling females were considered the reference group. To determine changes in pathways across aging, pairwise comparisons of normalized pathway scores were performed between ages within each sex, and Benjamini-Hochberg FDR adjusted *p*-values of < 0.05 were considered statistically significant.

### 2.5 Statistical Analyses

Statistical analyses were conducted with IBM SPSS (Armonk, NY), GraphPad Prism (San Diego, CA), and nSolver Advanced Analysis (NanoString Technologies). Normal distribution and homogeneity of variance were assessed before proceeding to the appropriate parametric or non-parametric tests. Two- tailed independent-samples t-tests or Mann-Whitney U tests were used to compare two groups with one independent variable. One-way analysis of variance (ANOVA) or Kruskal-Wallis H tests were used for one dependent variable and ≥ 3 groups. Two-way ANOVA was used to compare the interaction between age and sex on dependent variables. Three-way repeated-measures ANOVA was used to compare three independent variables in which one variable repeated. In the case that sphericity was violated, statistical values are reported from the Greenhouse-Geisser correction. Chi-squared tests for independence were used for categorical data. Correlations between behavioral measures and pathway scores or gene expressions were assessed by calculating the Pearson correlation coefficient.

Familywise α was maintained at 0.05 with Bonferroni adjustments for all *post-hoc* tests except the determination of differentially expressed genes, which used the Benjamini-Hochberg FDR adjustment with an adjusted *p*-value threshold of 0.1. Non-parametric data are graphed as box and whisker plots, and parametric data are graphed as mean ± standard error. Full statistics for all analyses are reported in Tables S1, S3, S4, S5, and S6.

## 3. Results

### 3.1 Aging broadly impacts socioemotional behavior

Mice were taken through a battery of behavioral tests that measure different endophenotypes of socioemotional behavior at 4, 10, or 18 months of age (Figure 1A; for statistics, see Table S1). We first assessed open arm avoidance on the elevated plus maze as a measure of anxiety-like behavior. Two-way ANOVAs revealed main effects of age and sex on time spent in the open arms (Figure 1B) as well as main effects of age and sex on the percent of open arm entries (Figure 1C). These results indicate decreased open arm avoidance in females as compared to males and increased open arm avoidance across aging irrespective of sex. We next assessed locomotor activity and center avoidance in the open field test and observed no effects of age or sex on this assay (Figure 1D-E). These results demonstrate that aging does not grossly impact locomotor activity and that sex- and age-related differences in avoidance behavior are assay-specific.

**Figure 1.**
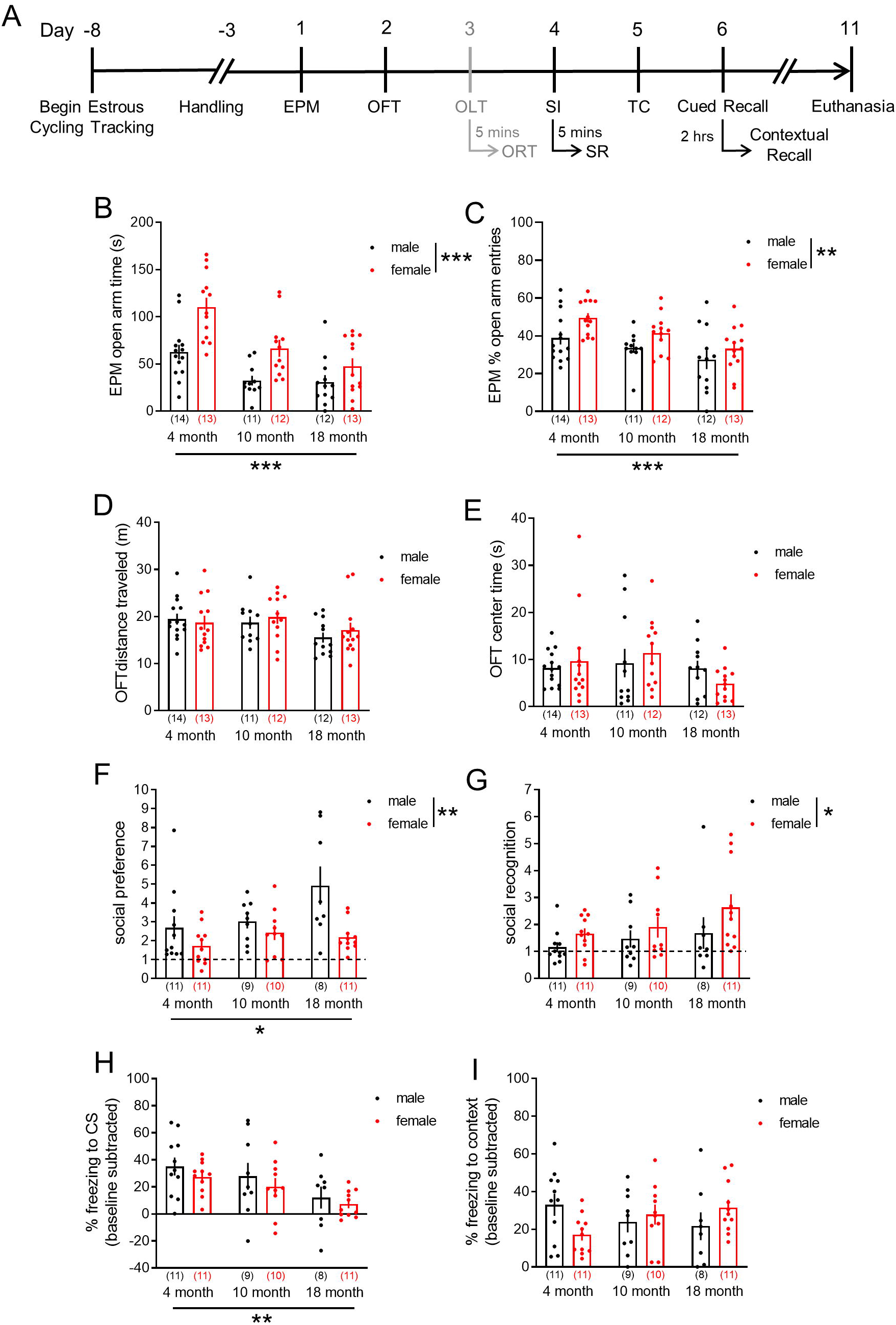
Sex- and aging-related differences in socioemotional behaviors. **A.** Experimental timeline. Male and female mice at 4, 10, or 18 months of age underwent a battery of socioemotional behavioral tests, and estrous cycle was tracked via vaginal (female) or sham (male) lavages for 8 days prior and throughout all experimental procedures. The object location test (OLT) and object recognition test (ORT) were performed but not presented. Five days following the last behavioral test, mice were euthanized. **B.** Open arm time on the elevated plus maze (EPM) was decreased with age independent of sex and was increased in females independent of age. **C.** Percent open arm entries on the EPM was decreased with age independent of sex and was increased in females independent of age. **D.** Distance traveled in the open field test (OFT) did not vary across age or between sexes. **E.** Center time in the OFT did not vary across age or between sexes. **F.** Social preference in the social interaction (SI) test was increased with age independent of sex and was decreased in females independent of age. **G.** Social recognition (SR) did not vary across age but was increased in females independent of age. Dashed lines in F-G represent no social preference/recognition. **H.** Normalized percent time freezing to the conditioned stimulus (CS) decreased with age during the cued threat memory recall test. **I.** Normalized percent time freezing in the conditioning context did not differ across age or sex. B-I, two- way ANOVA followed by Bonferroni-corrected *post-hoc* comparisons in the case of a significant interaction. For statistics, see Table S1. Asterisks below y-axes and to the right of legends depict significant main effects of age and sex, respectively. \**p* < 0.05, \*\**p* < 0.01, \*\*\**p* < 0.001. n/group denoted in parentheses under bar histograms.

To assess social behavior and cognition, mice next underwent the social interaction and social recognition tests with novel age- and sex-matched conspecifics. In the social interaction test, two-way ANOVA revealed a main effect of age and sex on social preference, with males displaying higher levels of social preference than females and levels increasing with age across both sexes (Figure 1F). In the social recognition test, two-way ANOVA revealed a main effect of sex, but not age, with females displaying higher levels of social recognition than males (Figure 1G). Together, these results indicate that preference for social interaction increases with age and that males exhibit increased social preference and decreased social recognition as compared to females.

Finally, emotional memory was assessed with auditory threat conditioning followed by cued and contextual memory recall the following day. Memory acquisition and recall was assessed by freezing, the dominant defensive behavioral response evoked by threatening stimuli in mice (Blanchard and Blanchard 1969, Fanselow 1980). To assess threat memory acquisition, we performed a three-way repeated-measures ANOVA to determine the impact of age and sex on freezing during the CS across the three CS-US trials during training (for descriptive statistics, see Table S1). We found a main effect of CS-US pairing, a main effect of age, and an interaction between age and CS-US pairing. To better understand this interaction, we performed pairwise comparisons between each age at each CS-US pairing, using the Bonferroni correction for multiple testing, and found increased freezing in 18-month compared to 4-month mice at the second CS, indicating accelerated memory acquisition in aged mice. Twenty-four hours later, cued threat memory recall was assessed by CS-freezing in a novel context.

Two hours later, contextual threat memory recall was assessed in the conditioning context. We first performed repeated-measures ANOVAs to assess within-group differences between pre-CS baseline freezing in the novel context, averaged freezing across the four CSs in the novel context, and freezing in the conditioning context (Figure S1). Young adult mice exhibited enhanced CS freezing compared to freezing in the novel and conditioning contexts, an effect that was lost with age. As we observed group differences in freezing not only during the pre-CS baseline period in the novel context but also during the pre-conditioning baseline period, we normalized our cued and contextual recall data by baseline subtraction to directly compare groups. Cued threat memory recall data was normalized by subtracting pre-CS baseline freezing from averaged CS freezing within each subject. Two-way ANOVA revealed a main effect of age with decreased normalized CS freezing as age increases (Figure 1H). Contextual recall data was normalized by subtracting pre-conditioning baseline freezing from contextual recall freezing. Two-way ANOVA revealed an interaction between age and sex on normalized contextual freezing (Figure 1I). To better understand this interaction, we conducted planned *post-hoc* comparisons of freezing between sexes at 4, 10, and 18 months, but no comparisons survived the Bonferroni correction for multiple testing. These results indicate that cued threat memory generalizes with age, as 18-month animals exhibit similar freezing during the pre-CS baseline period as during CS presentation in a novel context.

### 3.2 Sex differences in trajectory of aging-related changes in ventral hippocampal transcript expression

We next became interested in the ventral hippocampus, a brain region known to regulate socioemotional behaviors (Bannerman et al. 2004, Fanselow and Dong 2010) and that exhibits sex- and age-related differences (Wang et al. 2019, Williams et al. 2020, Porcher et al. 2021, Hodges et al. 2022). To better understand the impact of aging and sex on ventral hippocampal physiology, we performed transcriptional profiling of a subset of mice used for behavioral testing. RNA was extracted from ventral hippocampal tissue from mice at 4, 10, and 18 months of age and analyzed using the NanoString Neuropathology panel to quantify transcript counts from 760 genes (for list, see Table S2) with established involvement in neuropathological processes (Preuss et al. 2020, Cao et al. 2021). We performed pairwise comparisons to detect differentially expressed genes across age within each sex, setting FDR-adjusted *p* < 0.1 for detection of statistically significant changes (Figure 2).

**Figure 2.**
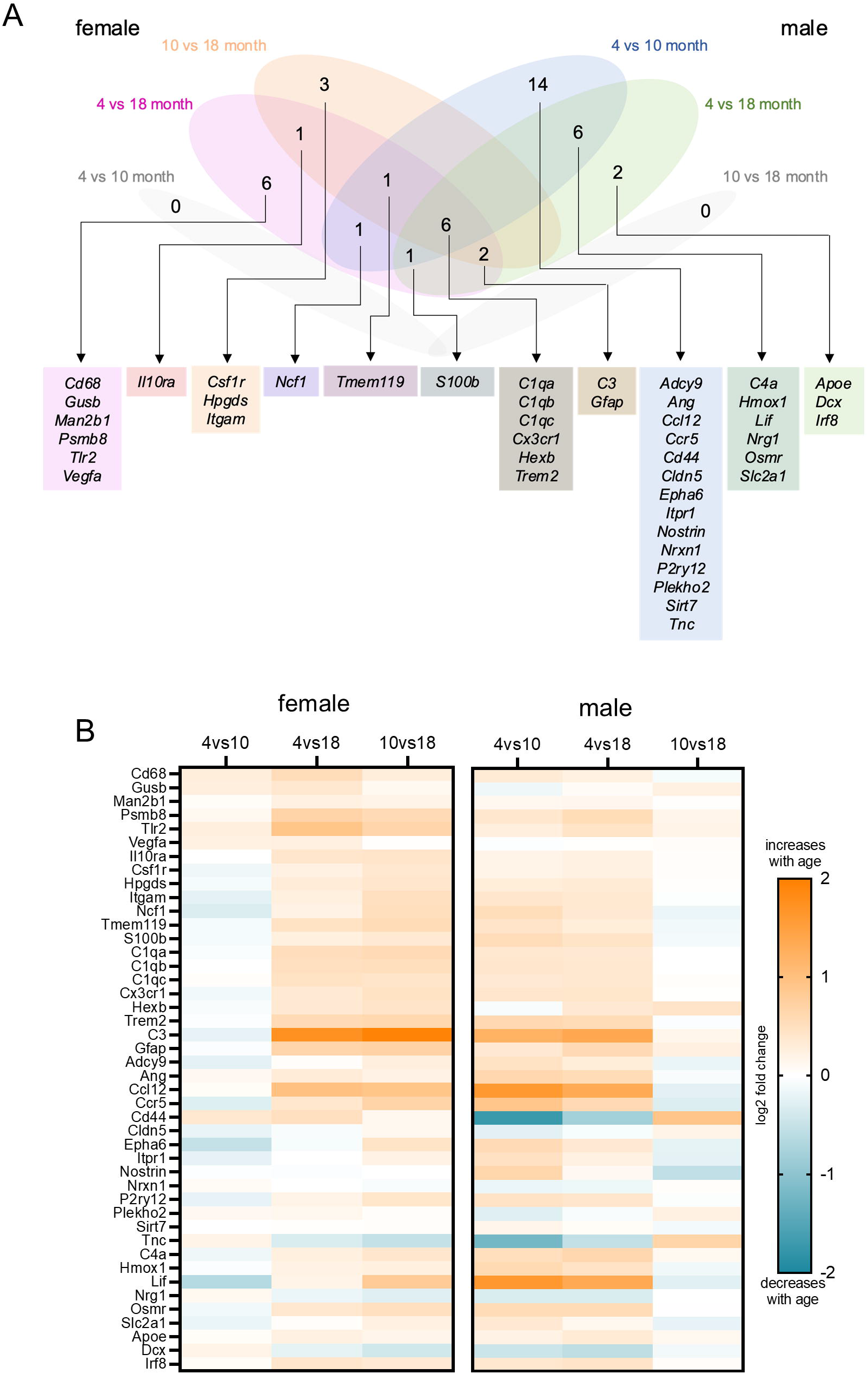
Sex differences in the timing and identity of transcriptional aging in the ventral hippocampus. RNA was extracted from the ventral hippocampus of male and female mice at 4, 10, and 18 months of age and analyzed using the NanoString Neuropathology Panel, which produced transcript counts for a list of 760 predetermined genes. **A.** Venn diagram demonstrating overlap and segregation of the 44 transcripts found to be differentially expressed across aging in males and females (FDR-adjusted p-values of *p* < 0.1). **B.** Heatmaps of differentially expressed transcripts across aging in females (left) and males (right), represented as the log2 fold change between the indicated timepoints. n=8/group. For statistics, see Table S3.

A total of 44 transcripts were differentially expressed across aging (Figure 2A; for statistics, see Table S3). Interestingly, the age of these transcriptional changes differed between the sexes with females being impacted later than males. In females, no changes were observed between 4 and 10 months of age. Eighteen transcripts were differentially expressed between 4 and 18 months, and 13 transcripts were differentially expressed between 10 and 18 months. In males, on the other hand, all age-related transcriptional changes occurred by 10 months of age, with no changes between 10 and 18 months. Seventeen transcripts were differentially expressed between 4 and 18 months, and 29 transcripts were differentially expressed between 4 and 10 months. Among the 44 total transcripts exhibiting age-related changes, only 11 were shared between males and females. We also found sex differences in the direction of differentially expressed transcripts (Figure 2B): transcript expression increased with age in females, whereas transcripts were both up- and downregulated in males. To directly compare the interaction between age and sex, we conducted two-way ANOVAs on these 44 differentially expressed genes (Figure S2; for statistics, see Table S4). In the case of significant interactions, Bonferroni-corrected *post-hoc* tests revealed more sex differences occurring at 10 months than at 4 and 18 months combined, further highlighting sex differences in the timing of transcriptional aging of the ventral hippocampus. Together, results indicate specific patterns in ventral hippocampal aging in which sex differences occur in timing, direction, and identity of transcriptional changes.

To determine the functional significance of age-related transcriptional changes, we next investigated changes in functional pathways using the nSolver Pathway analysis tool (for a list of transcripts included in each pathway, see Table S2). We performed pairwise comparisons of the 23 pathway scores and set statistical significance as FDR-adjusted *p* < 0.05 (Figure 3A; for statistics, see Table S5). We found that angiogenesis and autophagy increased with age in both sexes, whereas cytokines and neuronal cytoskeleton increased with age selectively in females.

**Figure 3.**
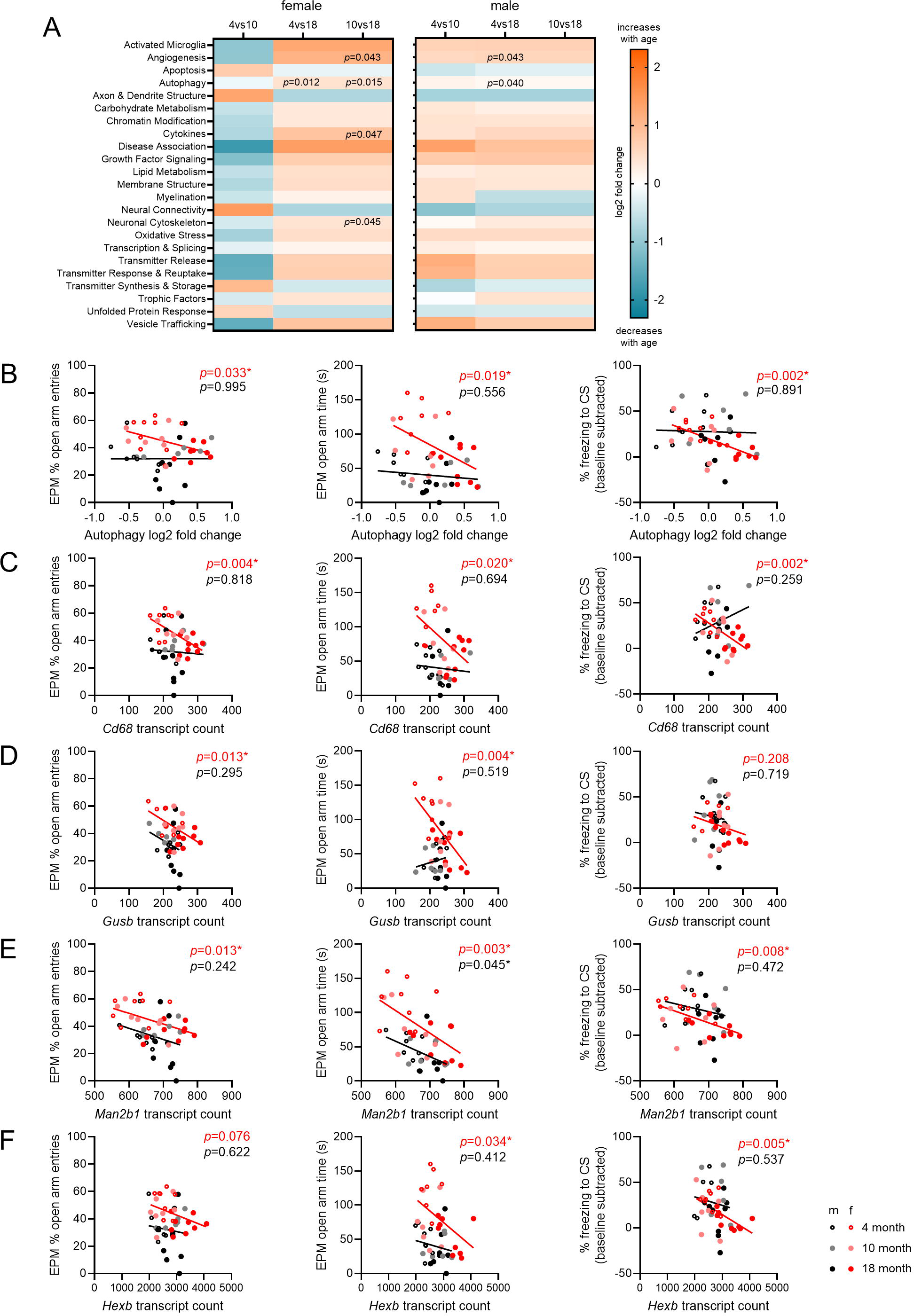
Sex-specific relationship between autophagy genes and socioemotional behaviors across aging. **A.** Transcript data were grouped by putative gene function using the nSolver Pathway Analysis Tool, revealing age-related changes in 4 of the 23 pathways (indicated with superimposed FDR-adjusted *p*-values of < 0.05). Subsequent analyses focused on autophagy as a pathway shared between sexes at the 4- versus 18-month comparison. **B.** The autophagy pathway score correlated with elevated plus maze (EPM) percent open arm entries (left), EPM open arm time (middle), and normalized freezing to the conditioned stimulus (CS) during cued threat memory recall (right) in females but not males. Subsequent analysis focused on the individual autophagy genes found to be differentially expressed across aging. **D.** *Cd68* expression was correlated with EPM percent open arm entries (left), EPM open arm time (middle) and normalized CS-freezing during cued recall (right) in females. **E.** *Gusb* expression was correlated with EPM percent open arm entries (left) and EPM open arm time (middle), but not normalized CS-freezing during cued recall (right), in females. **F.** *Man2b1* expression was correlated with EPM percent open arm entries (left) and normalized CS-freezing during cued recall (right) in females and with EPM open arm time (middle) in both sexes. **G.** *Hexb* expression was correlated with EPM open arm time (middle) and normalized CS-freezing during cued recall (right) in females. Legend in the bottom right corner applies to all graphs in B-F. B-F, Pearson’s correlations. For statistics, see Tables S5 and S6. \**p* < 0.05. n=8/group.

To identify differentially expressed genes that may be involved in the observed behavioral differences, we performed correlational analyses comparing each of the pathways to each of our behavioral measures (Figure 1) in order to determine if any pathway had a particularly strong correlation with socioemotional behavior changes across aging. Although several pathways showed significant age-related changes across aging or significant correlations with behavioral measures (Table S6), we chose to narrow our focus to the autophagy pathway because it uniquely demonstrated significant changes between the 4- and 18-month time points in both males and females, and it showed significant correlations with multiple socioemotional behavioral measures (Figure 3A-B). In females, autophagy pathway scores were negatively correlated with percent of open arm entries on the EPM, time spent in the open arms on the EPM, and the normalized percent of time spent freezing in response to the CS during cued threat memory recall. In males, however, autophagy pathways scores did not correlate with any behavioral measure (Table S6).

Due to this sex difference in the relationship between autophagy pathway scores and socioemotional behaviors, we chose to investigate the relationship between differentially expressed autophagy genes and behavioral measures. Of the four autophagy transcripts that were differentially expressed across aging (Figure 2), three were specifically increased in females (*Cd68, Gusb, Man2b1*) while one was similarly increased in both sexes across aging (*Hexb*). We then tested the correlations between these four genes and the three behavioral measures that correlated with autophagy pathway scores (Figure 3C-F; for statistics, see Table S6). We found that percent of open arm entries on the EPM was negatively correlated with *Cd68*, *Gusb*, and *Man2b1* in females, but not males. The amount of time spent in the open arms on the EPM was negatively correlated with *Cd68*, *Gusb*, and *Hexb* in females, and with *Man2b1* in both females and males. The normalized percent of time spent freezing to the CS during cued recall was negatively correlated with *Cd68*, *Man2b1*, and *Hexb* in females, but not males. Together, these results indicate that sex differences in trajectory of ventral hippocampal aging coincides with sex-specific transcriptional associations with socioemotional behaviors.

### 3.3 No impact of female reproductive status on socioemotional behavior or ventral hippocampal transcript expression

Because of the well-established influence of ovarian hormones on socioemotional behaviors and hippocampal function (Walf and Frye 2006), we next sought to determine the impact of reproductive senescence on socioemotional behaviors and ventral hippocampal transcript expression. To establish ovarian reproductive status, we categorized estrous cycle regularity in females across the three ages. The proportion of females exhibiting regular estrous cycles decreased between 4 and 18 months and between 10 and 18 months (Figure 4A-B). Complete arrest of the estrous cycle, an indication of reproductive senescence (Felicio et al. 1984), was observed for approximately half of the 18-month group, so we compared cycling and non-cycling females at this age. Consistent with decreased circulating estradiol levels (Evans et al. 1941), uterine index was decreased in the non-cycling group as compared to the cycling group (Figure 4C). Notably, this effect was not due to differed body weight between groups (Figure 4D). For socioemotional behavior, we observed no effect of cycling status on any outcome (Figure 4E-J). Finally, ventral hippocampal transcript analysis revealed no differentially expressed genes (Figure 4K) or changes in transcriptional pathways (Figure 4L) meeting the threshold of statistical significance in cycling versus non-cycling females at 18 months. Although these analyses are likely underpowered to detect subtle differences between cycling and non-cycling females, our findings suggest that reproductive senescence in aged females does not substantially impact socioemotional behavior or ventral hippocampal transcription.

**Figure 4.**
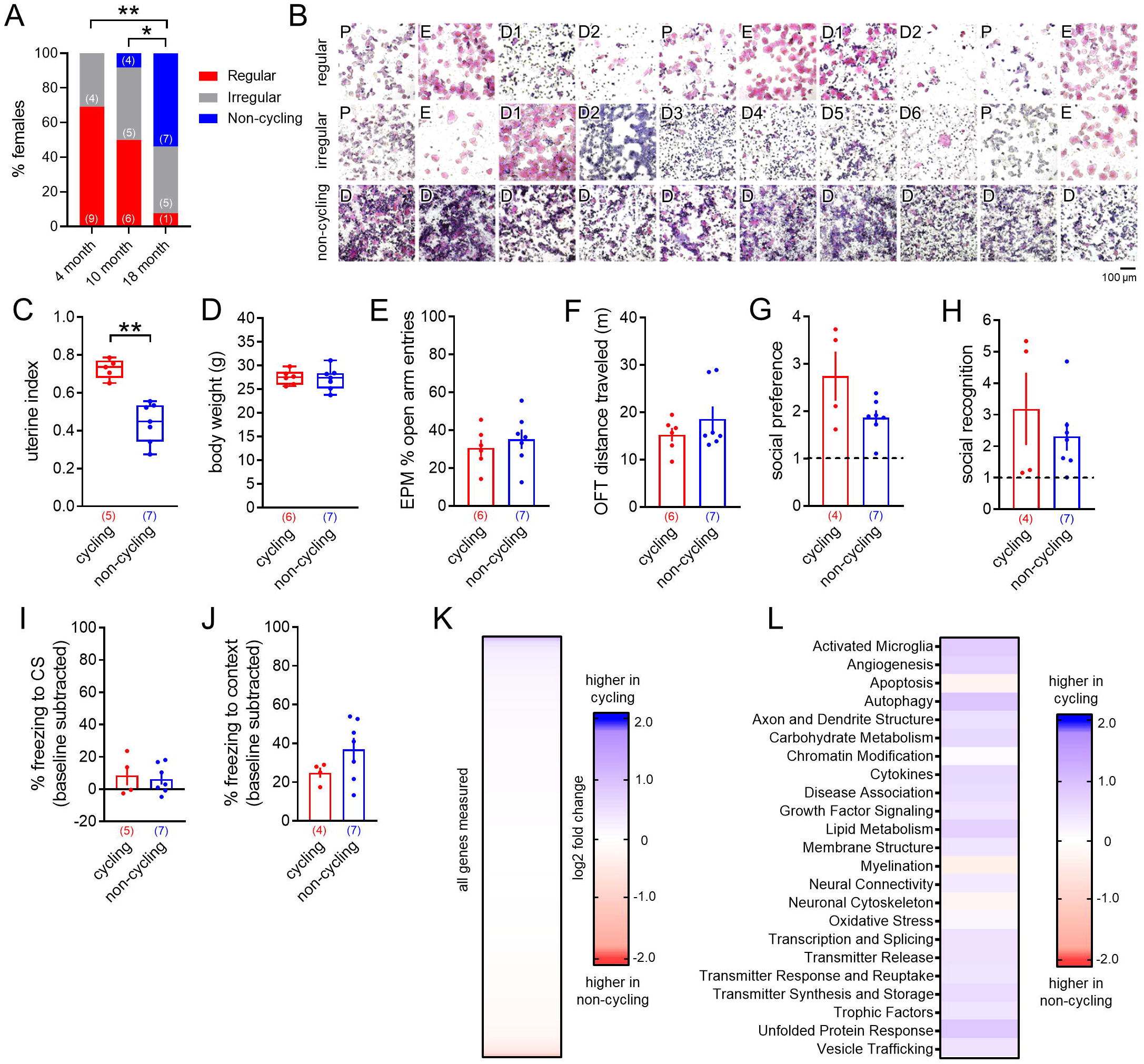
No differences in socioemotional behavior or ventral hippocampal gene expression between cycling versus non-cycling aged females. **A.** The proportion of females exhibiting regular estrous cycles was decreased between the 4- and 18-month and 10- and 18-month time points. **B.** Representative images of 10 consecutive days of vaginal cytology samples from regular, irregular, and non-cycling females. Scale bar (100 μm) is representative of all images. P, proestrus; E, estrus; D, diestrus. **C.** Uterine index was decreased in non-cycling versus cycling females. **D.** Body weight was not different between groups. **E.** EPM percent open arm entries was not different between groups. **F.** Open field test (OFT) distance traveled was not different between groups. **G.** Social preference was not different between groups. **H.** Social recognition was not different between groups. Dashed lines in F-G represent no social preference/recognition. **I.** Normalized freezing to the conditioned stimulus (CS) during cued threat memory recall was not different between groups. **J.** Normalized freezing to the conditioning context during contextual threat memory recall was not different between groups. **K.** Heatmap of expression of the 760 assayed transcripts (for ordered list, see Table S3). No differences were detected between cycling and non-cycling 18-month-old females. **L.** Heatmap of nSolver pathway scores. No differences were detected between groups. A, χ^2^ test for independence; C-D, Mann-Whitney U test; E-J, two-tailed t-test. For statistics, see Tables S1, S3, and S5. \**p*<0.05, \*\**p* < 0.01. n/group denoted in parentheses under bar histograms in A,C-K; n=4/group in K-L.

## 4. Discussion

This project provides the most comprehensive assessment of sex differences in socioemotional behaviors and novel sex-specific changes in the ventral hippocampal transcriptome across aging in young adult (4-month), middle-aged (10-month), and aged (18-month) C57Bl/6J mice to date. We report age-related differences in anxiety-like behavior, social preference, and threat memory generalization, as well as sex differences in anxiety-like behavior, social preference, and social recognition. These sex- and age-related behavioral changes were accompanied by sex-specific patterns of aging in the ventral hippocampus, with critical sex differences in both timing and direction of transcriptional changes, independent of reproductive senescence in aged females. Our results indicate sex differences in the trajectory of ventral hippocampal aging that may contribute to age- and sex-related changes in socioemotional behaviors.

Our findings indicate sex differences in anxiety-like behavior that persist throughout aging. As previously reviewed (Kokras and Dalla 2014), decades of work has shown that female rodents exhibit decreased anxiety-like behavior as compared to males across most behavioral tests, and that these effects may be driven by ovarian hormone levels across the rodent estrous cycle (Rocks et al. 2022). Here we report increased open-arm avoidance on the EPM in male versus female C57Bl/6J mice across all ages. Consistent with some (Darwish et al. 2001, Boguszewski and Zagrodzka 2002, Narita et al. 2006, Turner et al. 2012, Stanojlovic et al. 2019, Li et al. 2020, Hirano et al. 2021, Yanai and Endo 2021), but not all (Frick et al. 2000, Shoji et al. 2016, Shoji and Miyakawa 2019) previous work, we also report increased open-arm avoidance across aging. Notably these differences cannot be attributed simply to locomotor differences with aging, as we found no differences in distance traveled in the open field test. Our results therefore indicate that although there is a general age-related increase in anxiety-like behavior, sex differences in these behaviors persist across aging.

Contrary to some (Boguszewski and Zagrodzka 2002, Salchner et al. 2004, Hunt et al. 2011, Perkins et al. 2016, Shoji et al. 2016, Gerasimenko et al. 2020), but not all (Guan and Dluzen 1994, Shoji and Miyakawa 2019), previous reports in rodents, we found an age-related increase in social preference in C57Bl/6J mice. Interestingly, we further report opposing sex differences on social preference and social recognition independent of aging, with males displaying higher social preference, but lower social recognition, as compared to females. Both social preference and social recognition have shown to be profoundly impacted by gonadal hormone signaling (Choleris et al. 2009), yet how those impacts change with aging were undetermined. Here, our findings indicate baseline sex differences in social behavior that persist into old age, a time period where social behavior is known to dramatically improve cognitive and physical health outcomes (Seeman et al. 2001).

Our findings also demonstrate effects of age, but not sex, on cued threat memory dynamics. As previously reviewed (Bauer 2023), reports of sex differences in either cued or contextual threat memory processes vary greatly. Here we find that cued threat memory generalizes with age, as aged mice no longer exhibit enhanced freezing to the CS compared to the pre-CS baseline period in the novel context. Some (Liu et al. 2003, Feiro and Gould 2005, Gemma et al. 2005, Gould and Feiro 2005, Peleg et al. 2010, Shoji and Miyakawa 2019), but not all (Doyere et al. 2000, Blank et al. 2003, Villeda et al. 2011), previous reports have shown decreased cued memory recall across aging. However, these effects are difficult to disentangle from the age-related increase in threat memory generalization demonstrated here and elsewhere (Feiro and Gould 2005, Shoji et al. 2016, Yanai and Endo 2021), in addition to previously reported age-related deficits in context discrimination (Corcoran et al. 2002, Hernandez et al. 2022). Contrary to some (Stoehr and Wenk 1995, Oler and Markus 1998, Doyere et al. 2000, Corcoran et al. 2002, Gemma et al. 2005, Moyer and Brown 2006, Fukushima et al. 2008, Kaczorowski and Disterhoft 2009, Villeda et al. 2011, Ehlers et al. 2020, Yanai and Endo 2021, Hernandez et al. 2022) but not all (Gould and Feiro 2005, Aziz et al. 2019, Shoji and Miyakawa 2019) previous reports, we find no effects of age on contextual threat memory. However, in our study, context was conditioned in the background rather than foreground (Huckleberry et al. 2016). Future experiments should further test the perimeters of age-related threat memory generalization through both background cue discrimination as well as foreground contextual discrimination paradigms.

Although the hippocampus as a whole has long been studied in the contexts of both sex differences (Koss and Frick 2017) and aging (Rosenzweig and Barnes 2003), far less is known about the impacts of sex and aging on the ventral hippocampus specifically. Here, we provide the most comprehensive assessment of the ventral hippocampal transcriptome in males and females across aging to date. Despite the broad changes in socioemotional behavior we observed across aging, our analyses of ventral hippocampal transcripts surprisingly revealed only 44 (out of 760) differentially expressed genes across aging. Of those genes, 11 were similarly upregulated in both sexes across aging. These shared differentially expressed genes were primarily markers of microglial activation (*C1qa, C1qb, C1qc, C3, Cx3cr1, Ncf1, Tmem119, Trem2*) and angiogenesis (*C1qa, C1qb, C1qc, C3, Cx3cr1*), but also included genes associated with autophagy (*Hexb*), cytoskeleton (*Gfap*), and calcium signaling (*S100b*). All of these 11 genes have previously been shown to increase with aging in brains of male mice (Matarin et al. 2015, Ederer et al. 2022), and all but two (*C1qb*, *C1qc*) specifically in the hippocampus (Matarin et al. 2015, Mangold et al. 2017, Ederer et al. 2022, Lu et al. 2022). One study in female mice found aging-related increases in several of these genes (*C1qa, C1qb, Gfap, Hexb Tmem119, Trem2*) (Mangold et al. 2017), but until now a thorough comparison of sex differences in ventral hippocampal transcriptional changes with aging has been lacking.

In females, most differentially expressed genes were markers of activated microglia (*Cd68*, *Csf1r*, *Gusb*, *Psmb8*, *Tlr2*) and cytokines (*Csf1r*, *Il10ra*, *Vegfa*), though we also report female-specific upregulation of several genes associated with autophagy (*Cd68*, *Gusb*, *Man2b1*). Of these, *Cd68*, *Csf1r*, *Man2b1, Tlr2, and Vegfa* have been shown to increase in the aging hippocampus of female mice (Chen et al. 2018) and male mice in some (Matarin et al. 2015, Ederer et al. 2022), but not all (Wong et al. 2005, Yegla and Foster 2022), studies. Although aging-related changes have not been reported in brain levels of *Gusb*, sex differences across aging were recently reported in skeletal muscle tissue (Mishra et al. 2023). On the other hand, we report more variability in the functions of genes differentially expressed in males. We found notable representation of genes associated with angiogenesis (*Ang*, *C4a, Hmox1*, *Nrxn1*), axon and dendrite structure (*Adcy9*, *Apoe*, *Cldn5*, *Dcx, Nrg1*), cytokines (*Ccl12*, *Ccr5*, *Lif*, *Osmr*, *Plekho2*), and neural connectivity (*Ccl12, Itpr1, Nrxn1, Nrg1*) among male differentially expressed genes. Hippocampal expression of all except three (*Ccl12, Ccr5, Itpr1*) of these male-specific differentially expressed genes have previously been shown to change in males across aging in either mice (Matarin et al. 2015, Mangold et al. 2017, Lu et al. 2022) or humans (Berchtold et al. 2008), although we report opposite direction of changes for two genes (*Adcy9, Nrg1*). Additionally, *Dcx*—which we report is downregulated in the ventral hippocampus of aging males only—is considered a marker of adult neurogenesis (Couillard-Despres et al. 2005). We also find increased *Dcx* expression in males compared to females at 4 months (Figure S2), in accordance with a recent report demonstrating higher levels of neurogenesis in the ventral hippocampus of males versus females in young adulthood (Hodges et al. 2022), suggesting that *Dcx* may be a male-specific marker of the aging ventral hippocampus. Interestingly, we report similarly increased markers of cytokines in both sexes, though the specific genes involved differed between males (*Ccl12, Ccr5, Lif, Osmr, Plekho2*) and females (*Csf1r, Il10ra, Vegfa*). Previous reports in rodents (Mangold et al. 2017, Porcher et al. 2021) and in humans (Berchtold et al. 2008) demonstrate more dramatic aging-related increases in transcription of inflammatory and microglia-related genes in the brains of females versus males. Future work should explore whether these genes are therefore good candidates for markers of sex differences in neuroinflammatory responses with aging.

Notably, we report sex differences in both the timing and direction of transcriptional changes in the ventral hippocampus across aging. All female differentially expressed genes were upregulated with aging and occurred between the 10- and 18-month time points. On the other hand, males displayed a mix of up- and down-regulated genes that all occurred between the 4- and 10-month time points. These findings suggest accelerated aging in the ventral hippocampus of males compared to females, contrary to some (Yuan et al. 2012, Zhao et al. 2016), but not all (Berchtold et al. 2008), previous reports in the hippocampus and cortex of both rodents and humans, though the number of genes analyzed in these different studies varies greatly. Considering the transcriptionally unique identity of the ventral hippocampus (Dong et al. 2009, Floriou-Servou et al. 2018), our data may indicate sub-region and sex-specific trajectories of transcriptional aging within the hippocampus. Importantly, these differences do not appear to be mediated by circulating ovarian hormone levels, as we report no transcriptional differences between cycling versus non-cycling 18-month females. These negative effects are somewhat surprising considering recent work demonstrating changes in chromatin accessibility and transcriptional patterns across the estrous cycle in the ventral hippocampus of young adult mice (Jaric et al. 2019). However, the comparisons in the current study may be underpowered due to the low number of cycling females in the aged group. Aside from circulating hormone levels, other potential explanations for the seemingly delayed ventral hippocampal aging in females include X-chromosome linked resiliency (Davis et al. 2019). Future studies employing the four core genotypes mice would be required to resolve the influence of gonadal versus chromosomal sex on ventral hippocampal aging.

Together, our transcriptional data provide the most in-depth insight into sex differences in ventral hippocampal aging to date, enabled by the considerable sensitivity and broad scope of NanoString technology. However, one limitation of using the NanoString Neuropathology Panel is that our investigation is limited to genes specifically known to be involved in neuropathology. Considering the long history of sex bias in neuroscience research upon which this foundational knowledge is based (Beery and Zucker 2011), unbiased approaches such as RNA sequencing may therefore reveal more striking sex- and aging-related differences. Additionally, the false discovery rate correction (Benjamini et al. 2001) required by large datasets from NanoString technology may contribute to the lack of changes we report in certain genes (*Cd33, Ap2a2, Il6,* among others) shown to change with aging in other studies using lower throughput methods such as q-RT-PCR (Ederer et al. 2022).

Our initial goal in performing transcriptional analysis of the ventral hippocampus was to identify specific differentially expressed genes that may be regulating sex- and age-related changes in socioemotional behaviors. Comparisons of pathway scores across aging narrowed our focus to the autophagy pathway, which was uniquely increased in both sexes between the 4- and 18-month time points. Interestingly, autophagy pathway scores were negatively correlated with multiple measures of socioemotional behaviors in females, but not in males. When these same behaviors were analyzed in comparison to the four differentially expressed genes in the autophagy pathway, all were similarly correlated with socioemotional behaviors in females, with only one gene (*Man2b1*) also correlated with a behavioral measure in males. As autophagy dysfunction is heavily implicated in aging (Kaushik et al. 2021), these sex differences in the relationship between autophagy and socioemotional behaviors may represent key mechanisms underlying sex differences in ventral hippocampal aging. Both gonadal hormones and sex chromosomes have been shown to mediate sex differences in autophagy function (Shang et al. 2021), and it has been theorized that sex differences in autophagy may contribute to increased risk of neurodegenerative disorders in females (Congdon 2018). Considering the female bias we report in ventral hippocampal expression of autophagy-related genes with aging, these findings therefore suggest potentially novel female-specific mediators of socioemotional behavior across aging. Future studies should investigate causal links between these autophagy-related genes and socioemotional behaviors in order to better understand the mechanisms underlying these sex-specific associations.

In conclusion, this study provides the first broad assessment of sex differences in socioemotional behaviors across aging and suggests sex-specific trajectories of ventral hippocampal aging. Our data provide the most large-scale assessment of sex differences in the ventral hippocampal transcriptome across aging to date and are the first to consider the variables of age and sex in within-subject correlations between ventral hippocampal transcript levels and socioemotional behaviors.

These findings emphasize the importance of considering sex as a critical factor modulating the impacts of aging on socioemotional health and lay the foundation for future studies that may lead to therapeutic interventions targeted to women who make up a disproportionate percentage of the aging population.

## Acknowledgments

Many thanks are due to Averie Bunce and Amy Farthing for assistance with animal husbandry and vivisection, Arkady Bilenkin for assistance developing the RNA isolation protocol, Mariangela Martini for uterine dissection training, and Genna St. Amour for Nanostring training.

