## Supplemental figures for "Sex differences in socioemotional behavior and changes in ventral hippocampal transcription across aging in C57Bl/6J mice"

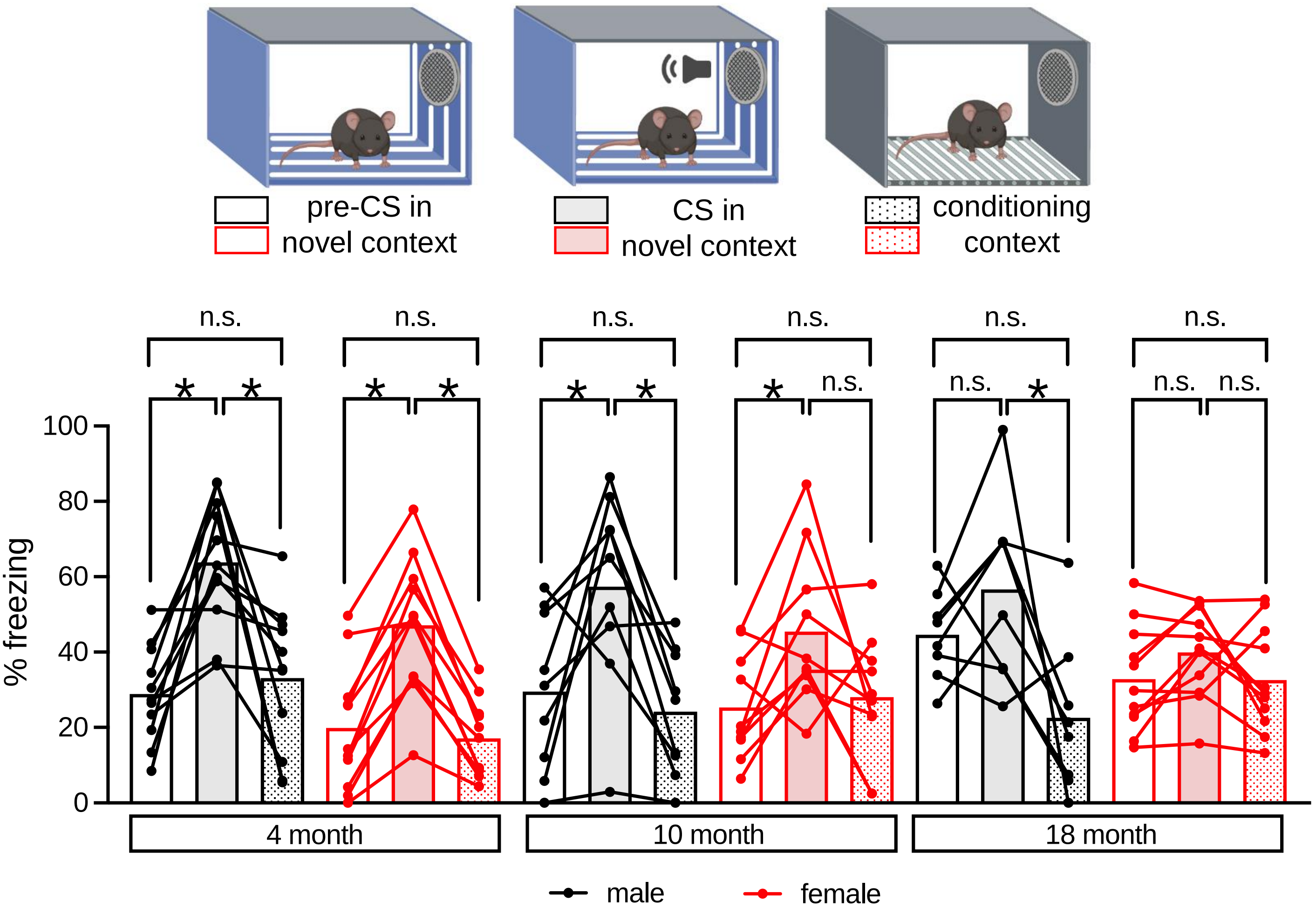


***Figure S1.*** *Cued threat memories generalize with age.* Comparison of within-group freezing during the pre-CS baseline period in a novel context (open bars), during the CS (filled bars), and during re-exposure to the conditioning context (dotted bars). For statistics, see Table S1. Repeated-measures ANOVA followed by Bonferroni-adjusted posthoc comparisons in the case of a significant interaction, * $p < 0.05$. Bars represent group means, and individual data points represent within-subject freezing across the three assays.

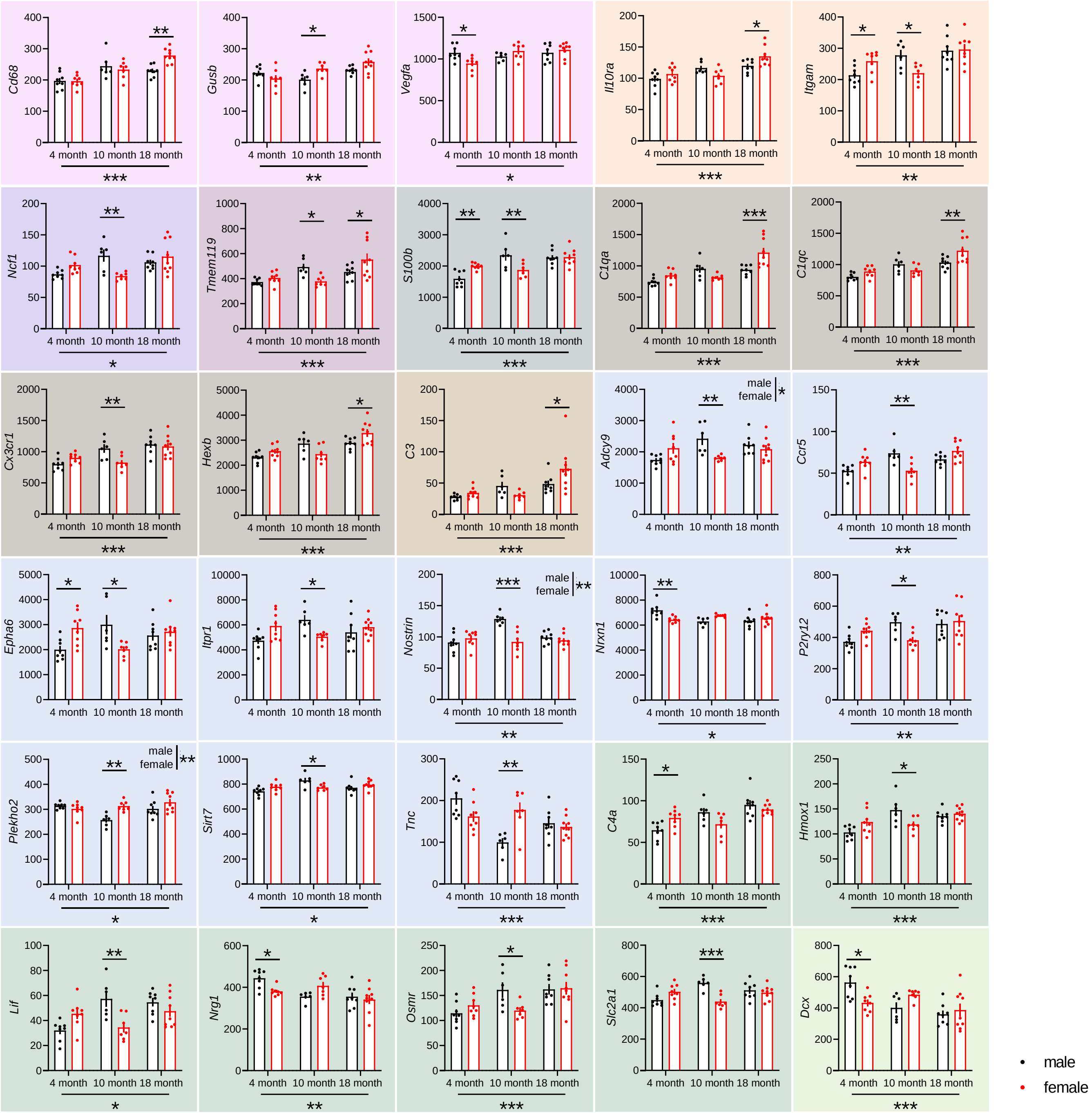


***Figure S2.*** *Differentially expressed genes displaying age by sex interactions.* Of the 44 differentially expressed genes, 30 displayed significant age by sex interactions with significant posthoc Bonferroni-corrected within-age sex comparisons (for statistics, see Table S4). Main effects of age are depicted with asterisks below the x-axes, and main effects of sex are indicated in the insets of *Adcy9* and *Nostrin*. Colors correspond to the groupings of differentially expressed genes depicted in Figure 2. Data presented as mean ± SEM. Two-way ANOVA followed by Bonferroni-corrected posthoc comparisons, *$p<0.05$, **$p<0.01$, ***$p<0.001$. n=8/group.
